# Meiosis in the human pathogen *Aspergillus fumigatus* has the highest known number of crossovers

**DOI:** 10.1101/2022.01.14.476329

**Authors:** Ben Auxier, Frank Becker, Reindert Nijland, Alfons J. M. Debets, Joost van den Heuvel, Eveline Snelders

## Abstract

Evidence from both population genetics and a laboratory sexual cycle indicate that sex is common in the fungus *Aspergillus fumigatus*. However, the impact of sexual reproduction has remained unclear. Here, we show that meiosis in *A. fumigatus* involves the highest known recombination rate, producing ~29 crossovers per chromosome. This represents the highest known crossover rate for any Eukaryotic species. We validate this recombination rate by mapping resistance to acriflavine, a common genetic marker. We further show that this recombination rate can produce the commonly encountered TR_34_/L98H azole-resistant *cyp*51A haplotype in each sexual event, facilitating its rapid and global spread. Understanding the consequences of this unparalleled crossover rate will not only enrich our genetic understanding of this emergent human pathogen, but of meiosis in general.

**One-Sentence Summary:** Genetic exchange between chromosomes during sex in *Aspergillus fumigatus* is higher than in any other known organism.

## Main Text

Invasive aspergillosis, caused by the ascomycete fungus *Aspergillus fumigatus* is a serious lifethreatening human disease. Clinical and environmental isolates are often resistant to antifungal azole treatments, developing from mutations either during clinical treatment or previous azole exposure in agricultural settings (*1*–*3*). Long suspected based on population genetic data, the sexual nature of *A. fumigatus* was confirmed by the discovery of a laboratory sexual cycle (*4, 5*). However, the impact of sex on genetic diversity remained unclear. The related genetic model *A. nidulans* has a high recombination rate, producing ~9-10 crossovers per chromosome (*6, 7*). With growing interest in population and mendelian genetics of *A. fumigatus* (*8–12*), understanding the outcome of sexual recombination is increasingly important. To investigate this, we constructed the first recombination map of *A. fumigatus*.

We used previously identified fertile strains AfIR964 and AfIR974 as parents in our cross (*13*). Genome assembly using short- and long-read data recovered all eight contiguous chromosomes for each parent (Table S1). Relative to the reference Af293, our parental genomes are largely syntenic, with little structural variation between them which could interfere with recombination (Fig. S1). After crossing, we randomly isolated 195 offspring from several cleistothecia, and generated ~90X depth of short-read data from each. Mapped against the AfIR974 parent, we identified 14,113 high confidence variants based on quality and segregation criteria (Table S2). These variants were unevenly spaced across the genome with a median inter-marker distance of 206 bp and a mean of 2010 bp.

Using a naïve criterion of recombination as non-parental adjacent markers (i.e.; A-B or B-A instead of A-A or B-B parental), we identified an average of 132.5 recombination events per offspring (16.6 per chromosome), producing a surprisingly long genetic map length of 14,843 cM (Fig. 1A; Table S3). The mean recombinant fraction between adjacent markers was 1% (max 38%), indicating that our markers can provide an accurate genetic map (Fig. S3). This criterion includes both crossovers, the physical exchange of homologous chromosomes, as well as smaller gene-conversion events that occur during resolution of meiotic double-stranded breaks (*14*). Since gene conversion events appear as two closely spaced crossover events, they can greatly inflate the genetic map (*14*). Therefore, this distance is an overestimate and requires correction. To remove gene conversions, we first used a rarefaction of our markers. Since gene conversion tracts are short, typically 0.5-2 kb (*15*), a high density-marker set has increased power to detect them while fewer markers are necessary to detect crossover events, which affect chromosomal segments orders of magnitude longer (*14*). Thus, increasing the gaps between markers, reducing the numbers of markers used, reduces the detection of gene conversions but less so for crossovers. Consistent with this, rarefaction of our markers reduced the map length asymptoticly (Fig. 1A). The reduced map lengths are not simply due to a reduced number of markers. For example, the 559 markers at 20 cM spacing resulted in a 12,512 cM map, while the 257 markers at 50 cM spacing produced a map of 11,966 cM — a ~50% reduction in markers only resulted in a 4.5% reduction in map length.

**Fig. 1.**
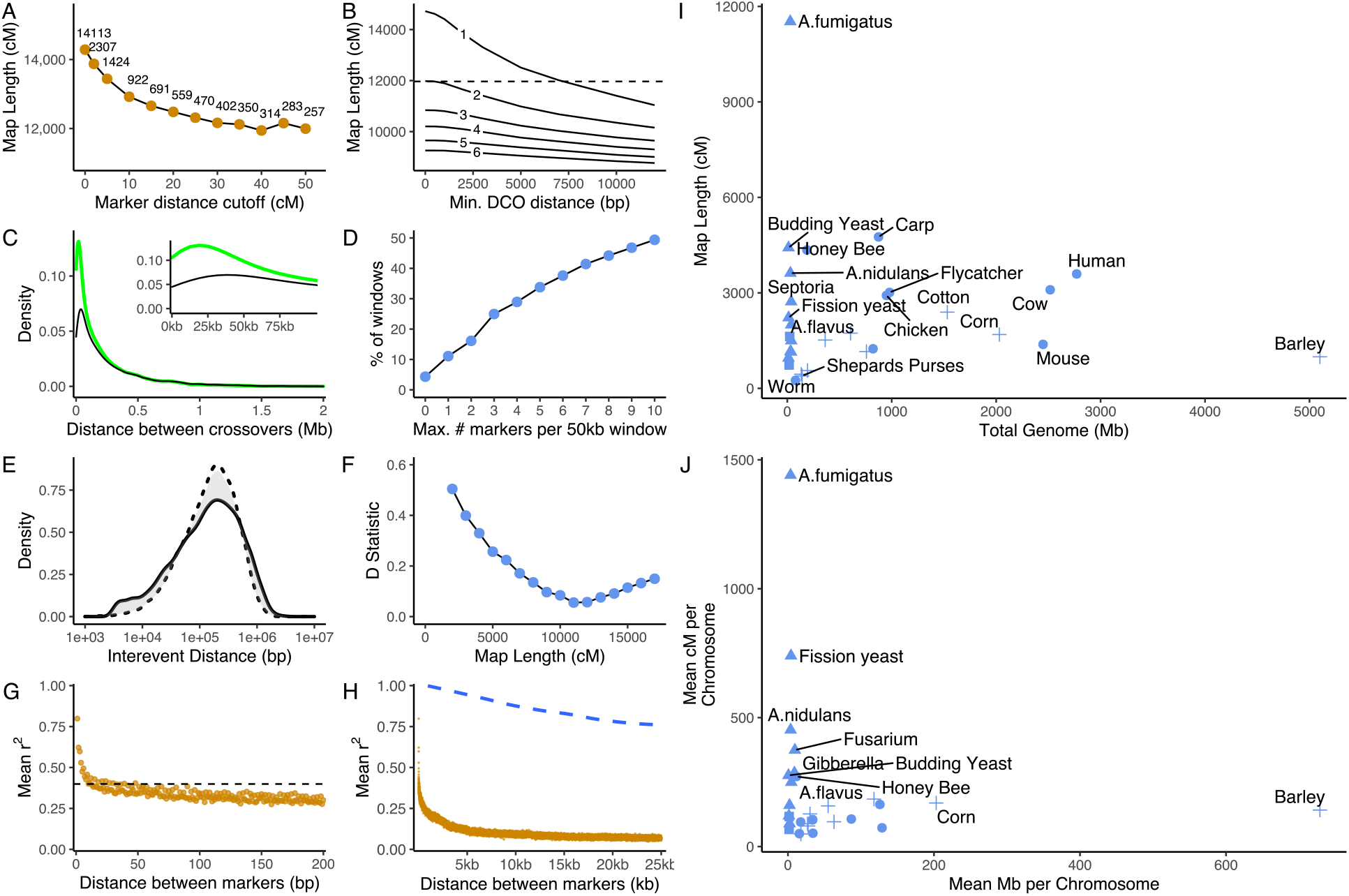
Genetic Map of *A. fumigatus*. **(A)** Total map length after removing markers closer than specified interval (x-axis), lengths represent mean of 5 samplings. Number of markers at specified spacing indicated near points. (**B**) Genetic map length after removing putative gene conversions based on length (x-axis). Criteria of minimum required markers is indicated in-line. **(C)** Distribution of DCO distances in raw data (green line) or after removing putative GC events (black line), inset shows zoom on first 100 kb distance. (**D**) Fraction of the 577 50 kb genomic windows with a minimum number of markers. **(E)** Distribution of DCO distances from empirical data (solid line) versus simulated data (dashed line) of an 11,000 cM genetic map. **(F)** Plot of D statistic (grey shading in E) comparing observed data to simulated genetic map lengths. (**G**) Decay of Linkage Disequilibrium (LD) across the *A. fumigatus* genome. Dots indicate average r^2^ between markers across 175 published individuals (*11*), dotted line indicates 50% reduction in LD. **(H)** LD decay across longer distances, orange dots indicate population level decay, while blue dotted line indicates LD of the mapping population. (**I**) Comparison of the genetic map of *A. fumigatus* to other species. **(J)** Similar to G but shown per chromosome. Data for I+J derives from (*16*) with the addition of (*6, 17*) and our data. Comparison to an alternate dataset is found in Fig. S4.

This rarefied distance is consistent with alternate methods. *Post hoc* criteria for removing gene conversions, such as minimum number of markers or length supporting a double crossover (DCO), had differing effects on the resulting map length (Fig. 1B). Increasing the minimum DCO length up to 12 kb resulted in an 11,037 cM map. However, increasing the minimum number of markers for a DCO had a stronger effect on map length, with a three-marker criterion reducing the map to 10,839 cM (Fig. 1B). This stronger effect of marker number is likely due to portions of the genome with few markers, as 16% of 50 kb genomic windows have less than three markers (Fig. 1D). As a further map length method, we simulated different map lengths of uniform crossovers across the genome. As the related *A. nidulans* lacks crossover interference, it produces uniformly distributed crossovers and the DCO distances form a Gamma distribution (*18*, *19*). We found that lengths between 11,000 and 13,000 cM best fit the data, minimizing the D statistic between observed and simulated data (Fig. 1E+F; additional map lengths Fig. S2).

We therefore conclude that the map length of *A. fumigatus* is between 11,000-13,000 cM, and we use the 11,966 cM estimate resulting from 50 cM marker rarefaction. To our knowledge this map length (0.422 cM/kb) is the longest estimated for any organism (Fig. 1I, Figure S4A) even after correcting for chromosome length (Fig. 1J, Fig. S4B). Further supporting this recombination rate, a previous study with different parents showed >5% recombination between two spore-color genes, *alb1* and *abr2,* spaced 8.3 Kb apart (>0.6 cM/kb) (*13*). However, the high recombination rate found in *A. fumigatus* does not appear to be widespread across *Aspergillus* species. High-density genetic maps for *A. nidulans* and *A. flavus* indicate lengths of 3,705 cM and between 1,500 and 2,000 cM, respectively (*6*, *17*). Using population level variation from a previously published dataset of 175 *A. fumigatus* German clinical and environmental isolates (*11*) the population level effect of sexual recombination is visible. The linkage between genetic variants decays rapidly, within 50 bp using the LD50 statistic (Fig. 1G+H). This rapid decay of linkage, with LD50 values for other species generally >1kb (*20*, *21*), indicates that sexual recombination in field populations is ongoing, as the single recombination events in our mapping population did appreciably decrease LD (Figure 1H dotted line).

As suspected based on the relationship to *A. nidulans,* we find no evidence of crossover interference – the nonrandom distribution of crossovers found in most organisms (*19*), although recombination is not uniform across the physical chromosome (Fig. 2B). We base this on three observations. First, as a direct measurement of interference (*22*), we find a coefficient of coincidence of ~1, indicating no deviation between observed and expected double crossover rates (Fig. 2C). Secondly, the distribution of crossovers per chromosome is very similar to the expected Poisson distribution (Fig. 2D). Finally, in the absence of interference the shape parameter of the interevent Gamma distribution, *v*, is expected to be 1, with values greater than 1 indicating interference (most species 5<*v*<30) (*19*). The *v*=0.74 (Fig 1C) of our cleaned dataset does not indicate interference. The related *A. nidulans* lacks crossover interference, as well as a meiotic synaptonemal complex (SC), similar to *Schizosaccharomyces pombe* (*18, 23*). We expect that *A. fumigatus* also lacks a SC, however sequence similarity of the structural proteins is only detectable within closely related species (*24*). Thus, bioinformatic analyses for presence/absence of SC proteins are not feasible as the nearest species characterized is *Sordaria* (*25*). Interestingly, the number of crossovers per individual in our data (min. 80; max. 300) was over-dispersed compared to the expected Poisson distribution (Fig. 2E). This is due to correlation between numbers of crossovers between chromosomes, a recently recognized phenomenon (Fig. 2F) (*26*, *27*).

**Fig. 2.**
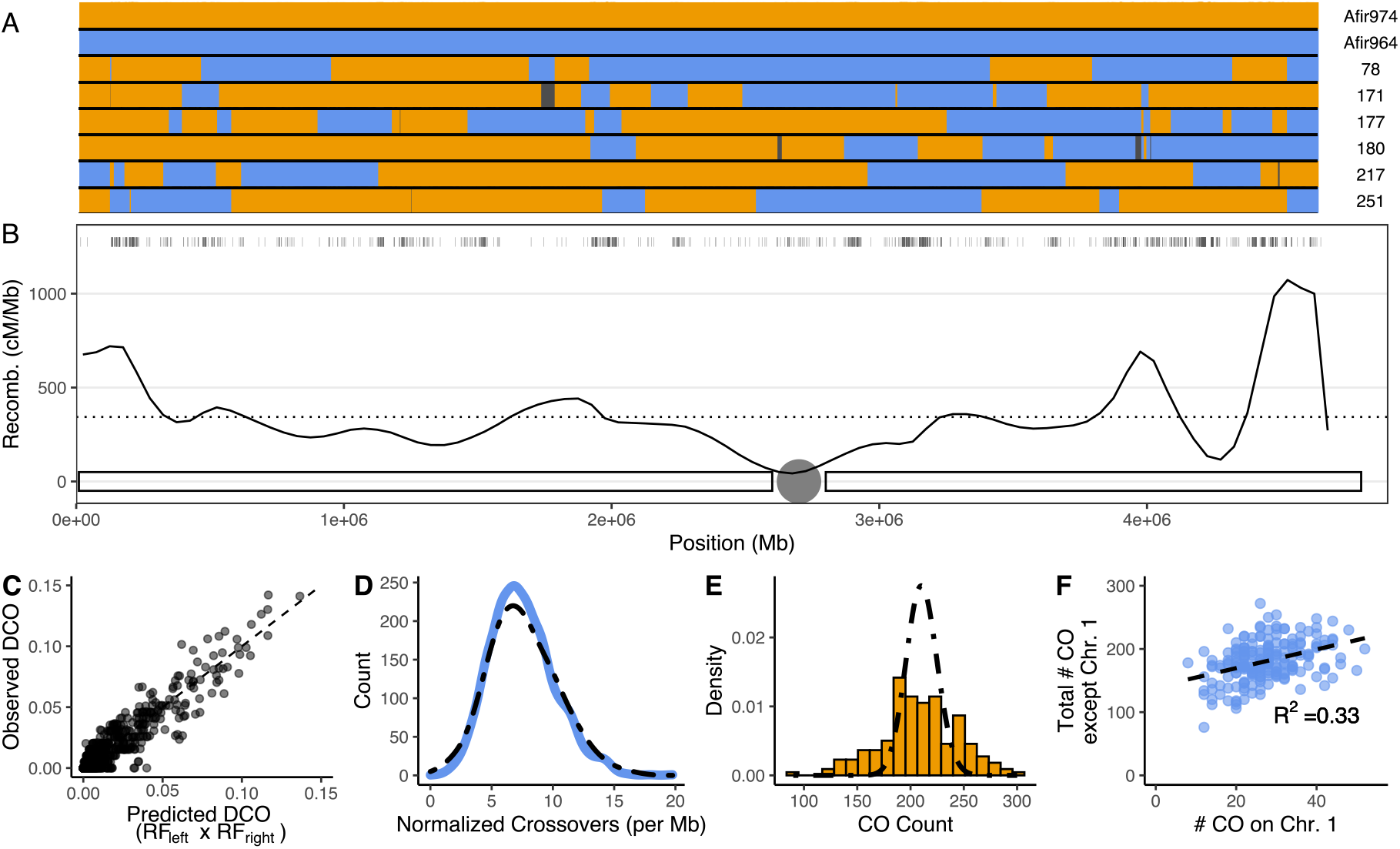
Crossover distribution in *A. fumigatus* shows no sign of interference. **(A)** Genotype parents and 5 representative offspring for Chromosome 1. Blue indicates Afir964 genotype, orange indicates Afir974. Black regions indicate regions removed from gene conversion criteria. **(B)** Recombination rate across Chromosome 1. Solid line indicates recombination rate across windows of 50kb, dotted line indicates mean recombination across Chromosome 1. Tick marks above indicate markers used for mapping, and chromosome with centromere is indicated below. Plots for all chromosomes are found in Fig. S5. **(C)** Correlation between predicted DCO based on adjacent intervals and observed DCO across the same interval in the 195 offspring. Diagonal dotted line indicates a 1:1 relationship, a coefficient of correlation of 1. **(D)** Histogram of normalized # of crossovers per Mb of chromosome, compared to Poisson distribution of same mean (dotted line) **(E)** Number of crossovers per individual in the dataset (orange bars) compared to the Poisson distribution of the same mean (dotted line) **(F)** Scatterplot of # of crossovers on Chromosome 1 versus the remaining total # of crossovers per genome, dots represent offspring.

As an independent line of evidence for this exceptional genetic map length, we tested how this translated to genetic mapping. Fortuitously, the parents had phenotypic variation in resistance to acriflavine (Fig. 3A), an antifungal with a rich history in fungal genetics (*28*). Phenotyping of the offspring on 50μg/mL acriflavine identified a single locus on chromosome 6 between positions 657 kb and 675 kb (Fig. 3B). The resolution of this genetic mapping, an 18 kb window, validates the high recombination rate as well as highlights the power of genetics in this species to identify novel mechanisms. The two variants in this region fall within the coding sequence of AFUA_6G03080 (Fig. 3B). This gene encodes an undescribed ABC efflux transporter in the family containing Multi Drug Resistance 1 (*mdr1*) (*29*). The difference in LOD scores between these two variants, from recombinant offspring across this distance of 410 bp, allows for immediate fine mapping of the presumed causal variant (PheàCys). While genetics has previously been used to identify resistance mechanisms in *A. fumigatus,* this study utilized sequentially evolved isogenic clinical isolates (*30*). Here we show that variants in *A. fumigatus* can be easily mapped with wild-type isolates, even for genes without known mutant phenotypes.

**Fig. 3:**
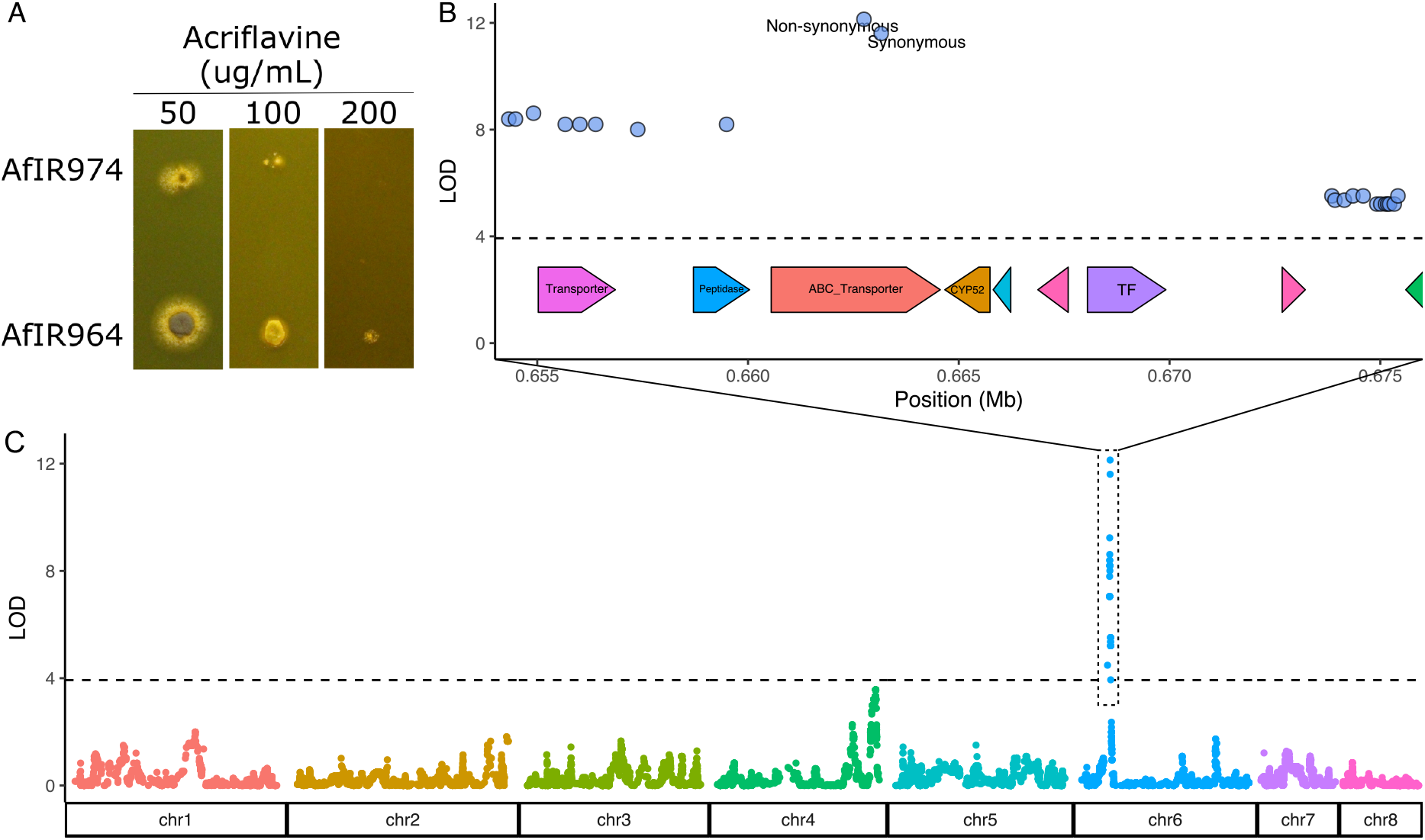
Validation of *A. fumigatus* recombination rate through mapping of acriflavine resistance. **(A)** Phenotypes of both parental strains grown at increasing concentrations of acriflavine after 2 days incubation **(B)** Closeup of the significant QTL window. Gene models shown below the dotted line indicate location of genes in the Afir964 parent. **(C)** Genome-wide significance values for acriflavine resistance in the cross between Afir964 and Afir974. Horizontal dotted line indicates the permutation-based significance threshold with α=0.05.

The high recombination rate of *A. fumigatus* may further explain observed patterns of antifungal resistance, a major concern in the clinical setting (*2*). Azole-resistant environmental isolates commonly contain haplotypes of 2-3 variants in the *cyp*51A gene involved in ergosterol synthesis. These resistant haplotypes are composed of a tandem repeat (often 34 or 46bp) in the promoter element (TR) combined with at least one non-synonymous polymorphism (e.g. TR_34_/L98H or TR_46_/Y121F/T289A). There are strong epistatic effects, as the effects are nonadditive between specific combinations of non-synonymous coding polymorphisms and promoter mutations causing increased antifungal resistance (*31, 32*). Curiously, it has been noted that in other fungal species resistance to the same azoles is instead due to single mutations in *either* the promoter *or* the coding sequence (*3*). The *cyp*51A haplotypes in *A. fumigatus* have been hypothesized to arise from mutation/tandem duplication at one position, and then an additional mutation/tandem duplication at the second position (*3*). Our data here suggests an alternative explanation of each mutation arising in independent strains, and then being united through recombination, which is considered a general benefit of sexual recombination (*33, 34*).

As our parental strains had no variation within or near the *cyp*51A gene, recombination could not be observed in this region, although it has been noticed at a population level (*11*). However, the 0.422 cM/kb recombination rate indicates that if one parent had the TR_34_ variant, and the other had the L98H variant (365bp apart), then 0.075% of offspring would have both resistance variants, (99.85 with either parental single variant and 0.075% with neither) (Fig. 4A). Recombination in the TR_46_ haplotype is expected to be slightly higher (TR_46_ and Y121F spaced 434bp apart). Since a single fruiting body produces >10,000 spores (*13*), recombinants within *cyp*51A gene are surprisingly, expected in each sexual event.

**Fig. 4:**
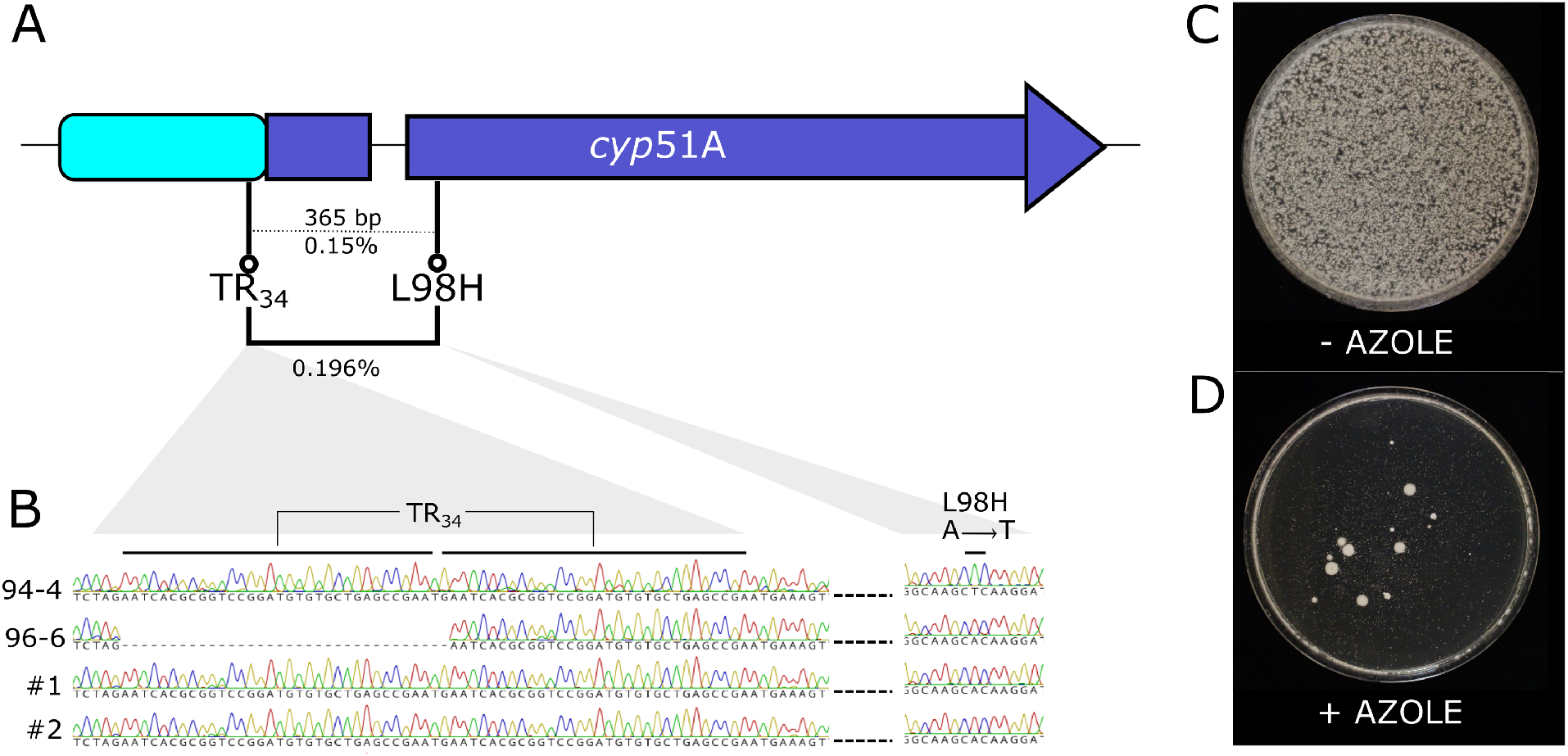
**Sexual recombination consistently** produces recombinant *cyp*51A haplotypes. **(A)** Diagram indicating distances and inferred recombination rates between positions in the *cyp*51A gene, leading to commonly encountered TR_34_ and TR_46_ azole resistance haplotypes. Base pair distances are indicated above dotted line, predicted recombination rates indicated below. Observed recombination rate from B indicated with solid line connecting TR_34_ and L98H. **(B)** Chromatograms of *cyp*51A sequence showing TR_34_ region and L98H region from parental strains with either variant, as well as two randomly selected azole resistant offspring **(C)** Ascospores of a single sexual fruiting body on non-selective media. **(D)** As above, but with addition of 10μg/mL itraconazole, which is above the tolerance of either single mutant strain. Large colonies indicate resistant offspring. Non-recombinant offspring are visible as scattered small dots. Note that plate for Fig. 4C was incubated 16 hours less than D to prevent overgrowth of abundant colonies. Images of all cleistothecia used are found in Fig. S6

To validate this predicted intragenic *cyp*51A recombination, we crossed previously generated single mutants (*35*) with AfIR974, generating sexually fertile offspring with either L98H or TR_34_ variants which each confer intermediate levels of azole resistance (4μg/mL itraconazole). After crossing these, we selected recombinants based on their positive epistatic effect by incubation on 10μg/mL itraconazole (Fig. 4C+D). These highly tolerant offspring were produced at a rate of 0.096% (mean number resistant colonies 18, mean total ascospores 18,708). Sequence analysis of the itraconazole resistant offspring confirmed them as recombinants, with both the L98H and TR_34_ variants (Fig. 4B).

Fungi are known to have higher recombination rates (*36*), but our finding of up to 64 crossovers per chromosome in *A. fumigatus* brings this to a new level. Across eukaryotes generally only a few crossovers occur per chromosome (*37*), and a negative correlation between population size and recombination rate indicates that excess crossovers are detrimental (*38*). This detriment can result from the breaking up of beneficial combinations of mutations (*33*) as well as physical meiotic defects, for example contributing to Trisomy 21 in humans (*39*), however this has been contested by recent experiments (*40*). This higher crossover number may be explained as the lack of a SC requires an increased number of crossovers to ensure at least one crossover per chromosome, necessary for proper meiotic chromosome reduction (*18, 23*). If this high crossover rate is simply a consequence of lacking a SC, it is unclear why the rate is so much higher than found in *A. nidulans* and *S. pombe,* which also lack a SC.

Our data shows that a physical limitation of crossover number does not explain the general low crossover rate across eukaryotes. Under a narrow range of parameters, a constantly changing environment can select for higher recombination rates (*41*). What ecological or evolutionary factors differentiate *A. fumigatus* from other *Aspergillus* species with a lower recombination rate is unknown. Whatever the evolutionary cause, this recombination rate likely facilitates azole-resistant haplotypes (*3*). This can now explain the rapid and global distribution of the azole resistant haplotypes, as sex appears to occur in bulb waste material heaps containing azole residues (*42*–*44*). This high recombination rate also affects the interpretation of population-level genome scans as it practically eliminates linkage between genes/markers (*12*). Understanding the adaptive advantage, or lack thereof, of this unparalleled number of crossovers requires further enquiry.

## Acknowledgments

We thank Dr. Eugene Gladyshev and Dr. Raphael Mercier for input on a previous version of this manuscript. We gratefully acknowledge Dr. W.M. Moye-Rowley for sharing the L98H and TR_34_ single mutants used in the crosses for Figure 4. We thank Dr. Paul Verweij and Dr. Paul Dyer for sharing the AfIR964 and AfIR974 parental strains. Dr. Erik Wijnker as well as other members of the Laboratory of Genetics provided valuable feedback during the writing process.

## Author contributions

Conceptualization: AJMD, ES

Data Curation: N/A

Formal Analysis: BA, JVDH

Funding acquisition: ES

Investigation: BA, ES, JVDH

Methodology: AJMD, BA, ES, FB, JVDH, RN

Project Administration: N/A

Resources: N/A

Software: BA, JVDH

Supervision: ES

Validation: N/A

Visualization: BA, JVDH

Writing – original draft: BA

Writing – review & editing: AJMD, BA, ES, JVDH

## Data and materials availability

All fungal strains used are freely available from the corresponding authors. Data and code supporting the analysis used are available at https://github.com/BenAuxier/aspergillus_recombination. Genome assemblies of parental strains AfIR964 and AfIR974 are undergoing submission as SUB10510768 and SUB10510426, respectively. Illumina data for all offspring is currently under submission at NCBI SRA SUB10935968.

## Supplementary Materials

Materials and Methods

Figs. S1 to S6

Tables S1 to S3

References (*45–62*)

### Sexual cross

Strains were crossed by plating both strains on Oatmeal Agar (72.5g/L; Difco Chemical), followed by incubation at 30°C for 4 weeks. Ascospores were harvested by manually isolating mature cleistothecia, and asexual conidia were removed by rolling in water agar. After crushing cleistothecia in water, the spores were heatshocked for 60 minutes at 70°C to kill remaining asexual conidia and hyphae. Aliquots of this spore suspension were then plated on Minimal Media (*45*) with 0.1% Triton X-100 to restrict colony size. Isolated single-spore colonies were transferred to MEA slants (30g/L Malt Extract; 1mg/L CuSO4) and incubated at 37°C.

### DNA isolation and sequencing

DNA was extracted using a modified method from (*46*) phenol-chloroform extraction from 24-hour old mycelial mats from both parents and all offspring. Conidia were added to 2mL of liquid Malt Extract media (30g/L Malt Extract), and incubated for 48 hours at 37°C. The resulting mycelial mat was removed to a microcentrifuge tube, to which 5-6 2mm glass beads were added. This tube was then frozen in liquid nitrogen, and homogenized with an Ivoclar Vivadent Silamat S6 homogenizer for 10 seconds at 4500 rpm, twice. To this mycelial powder, 530mL of Breaking Buffer (2% Triton X-100, 1% SDS, 10mM Tris-HCl pH 8.0, 1mM EDTA pH 8.0) and 20mL of 20mg/mL Proteinase K was added. After vortexing to mix, the samples were incubated for 1.5 hours at 56°C. Following this, 550mL of 24:24:1 Phenol:Chloroform:Isoamyl Alcohol was added, and samples were gently mixed on a rotary shaker for 10 minutes. Following this, the layers were separated using a table-top centrifuge at 14,000rpm for 15 minutes. The extraction step was repeated on the aqueous layer. DNA was precipitated from the aqueous layer using an equal volume of isopropanol, and washed using first 96% ethanol, followed by a wash with 70% ethanol. The resulting pellet was dried at room temperature and resuspended in 100mL of warm TE buffer (10mM Tris-HCl, 1mM EDTA, pH 8.0) and 1mL of RNAse I (Promega) was added. This purified DNA was incubated for 1 hour at 37°C to digest RNA, then frozen for storage. For each offspring and parental strain, an aliquot of DNA was processed for Illumina 150 bp paired-end sequencing (Novogene Inc.). For each of the parental strains, an additional DNA aliquot was then processed for the Oxford Nanopore platform by first using the Circulomics SRE kit according to manufacturer’s instruction to remove short DNA fragments, followed by the SQK-LSK-109 Genomic DNA by Ligation kit. Libraries for both parents were sequenced to ~100X depth using an Oxford Nanopore R9.4.1 flowcell (Supplemental Table 1). Basecalling was performed with Guppy software at high accuracy(3.2.8).

### Genome assembly

A hybrid assembly strategy was followed, merging an assembly using the minimap2 (-x ava-ont, 2.17-r954) / miniasm (0.3-r179) and canu (v.2.1.1, trim-assemble genomeSize=30m errorRate=0.05 -nanopore-raw) / racon (v1.3.1, -m 8 -x -6 -g -8 -w 500) pipelines (*47*–*50*). For the latter, bwa-mem mapping was used (v0.7.15, -x ont2d)(*51*). Both these assemblies were followed by pilon (v1.23) polishing before merging for which bwa-mem was used for mapping after which mapping was sorted and filtered (-q 20) using samtools (v1.9) (*52, 53*). After merging these assemblies using a custom script (Supplemental File, cutom.merge.sh), another round of racon correction and pilon polishing assured high contingency and accuracy. To test for genome completeness, BUSCO (v3.1.0) scores were calculated using the eurotiomycetes_odb9 database (--mode genome --species aspergillus_fumigatus) (*54*). The final genome was annotated with augustus (v2.5.5, --gff3=on --extrinsicCfgFile=extrinsic.E.cfg) using a hints file that was produced by using blat (v36.0, -t=dna -q=rna -minIdentity=90 -minMatch=1 -minScore=10) to search the Af293 reference genome coding sequences against the assembled genome (*55, 56*).

### Alignment, variant calling

Raw Illumina reads from both parents were aligned to the AfIR974 genome with bwa mem (v0.7.15), and subsequently filtered using samtools view (v1.9, -q 20), after which the resulting bam file was sorted. Reads groups were added using gatk AddOrReplaceReadGroups (v4.1.6.0)(*57*). Resulting bam files were used to call variants using freebayes (v0.9.21) (*58*). This raw VCF file was then annotated for effects of the variants using snpEff (v4.3t) using the above described augustus annotation.

### Offspring and Variant filtering

We performed a conservative variant filtering using vcfR (v1.12.0) (*59*). First filtered the raw vcf for read frequency (>0.95 between parents) and coverage (20 < coverage < 150 for both parents) to identify loci that were differentiated between parents. This allowed filtering for individuals that were either not offspring of the two parents, exactly similar to one of the parents, showed heterozygous allele frequencies or and lastly duplicate individuals. This filtering removed 52 individuals that were parental, indicating significant conidial survival despite 60 minutes incubation in a heating block at 70°C. This contradicts previous reports of conidial death at incubation at 70°C (*13*), likely due to the difference in heat transfer between a heating block with air gaps and a more effective water bath. We then removed loci that often showed a deviance from 0 or 1 in read frequency (those loci that on average deviated more than 1%), which removed 2.8% of the loci. Finally, genotypes were called per individual if a locus had a coverage higher than 15 and the alternative allele frequency of either AfIR974 or AfIR964 was 90% or higher.

Recombination between markers was determined based on physical ordering of said markers. After determining the excess genetic map length from gene conversions, excess recombination events were removed using the cleanGeno() function of R/qtl2.

### Population Linkage and Linkage Decay

To assess the impact of recombination in wild populations of *A. fumigatus,* we downloaded publicly available data from NCBI BioProject PRJNA697844. Of the 188 samples in total, 13 samples were removed as they are from the distantly related Cluster 1 as described in the original publication (*11*). Short reads of the remaining 175 individuals were then mapped to the reference Af293 genome using bwa-mem2 (*60*). Variants were called using freebayes as described above, and low quality sites as well as sites with excess missing data were filtered based on the following criteria “QUAL > 100 & AN > 290”. A further filter was applied to remove sites with 5 or more heterozygous calls. The result VCF file was processing using PLINK (v1.90) (*61*). LD was calculated using the r^2^ statistic, up to 25 kb of distance between sites, and results were aggregated using a previously published custom script (*12*). The VCF file used for the genetic mapping was processed similarly.

### Genetic mapping of Acriflavine Resistance

Acriflavine resistance was determined by plating ~10,000 spores in a 10uL droplet on Complete Media with 50μg/L acriflavine. Droplets were incubated for 3 days at 37°C before phenotyping. Growth/no growth was scored as a binary trait, and QTL mapping was performed with R/qtl2 using the scanone() to detect the responsible QTL (*62*). Genes were visualized using the gggenes package (v0.41) (https://wilkox.org/gggenes/), based on the GFF annotation of the AfIR974 parent.

### *cyp51A* Intragenic recombination

To assess recombination within *cyp51A* the strains SPF94 (AfS35 cyp51A^TR34^::*hph*) and SPF96 (AfS35 cyp51A^L98H^::*hph*) were each crossed with AfIR974 on Compost Agar media (60g/L Freeze Dried Mushroom Compost, 20g/L Agar). After incubation for 8 weeks, mature cleistothecia were harvested and heatshocked as above. Colonies resistant to both 4μg/mL itraconazole as well as 150μg/mL hygromycin (Ducha Chemicals) were isolated and mating type was determined using previously described primers (*5*). Offspring of compatible mating types with either TR_34_ or L98H variants were then crossed (94-4: TR_34_ MAT-1; 96-6: L98H MAT-2). From this cross single cleistothecia were isolated and heatshocked in 500uL H_2_O+0.05% Tween-80. From this ascospore suspension, 4 aliquots of 3μL were used for count plating on Malt Extract Agar (30g/L Malt Extract, 15g/L Agar), and the remained plated on large (16cm diameter) petri plates with Malt Extract Agar, either with or without 10μg/mL itraconazole. Count plates were incubated for 36 hours, and itraconazole plates were incubated 72 hours.

**Supplemental Fig. 1:**
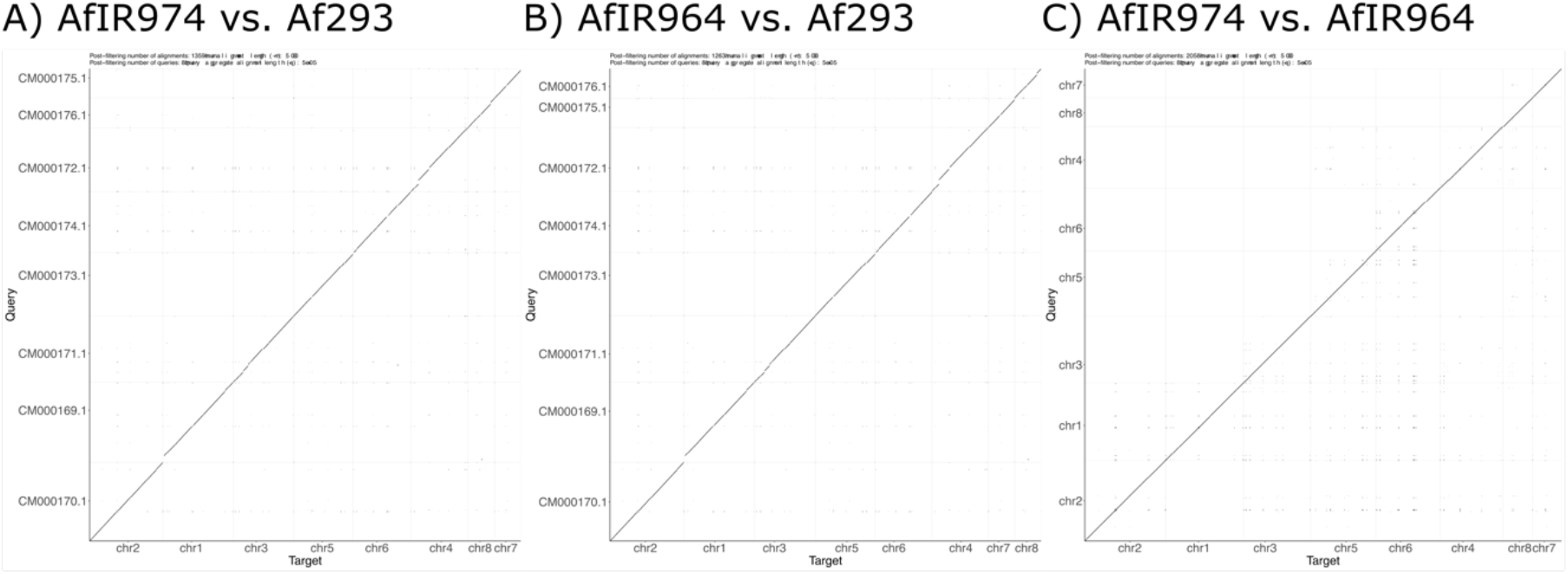
Synteny of two parental strains and reference strain Af293. **(A)** Dotplot comparison of sequence similarity using minimap2 of AfIR974 assembly against reference Af293 assembly. **(B)** Comparison of AfIR964 against Af293. **(C)** Comparison of two parental strains AfIR964 and AfIR974.

**Supplemental Fig. 2.**
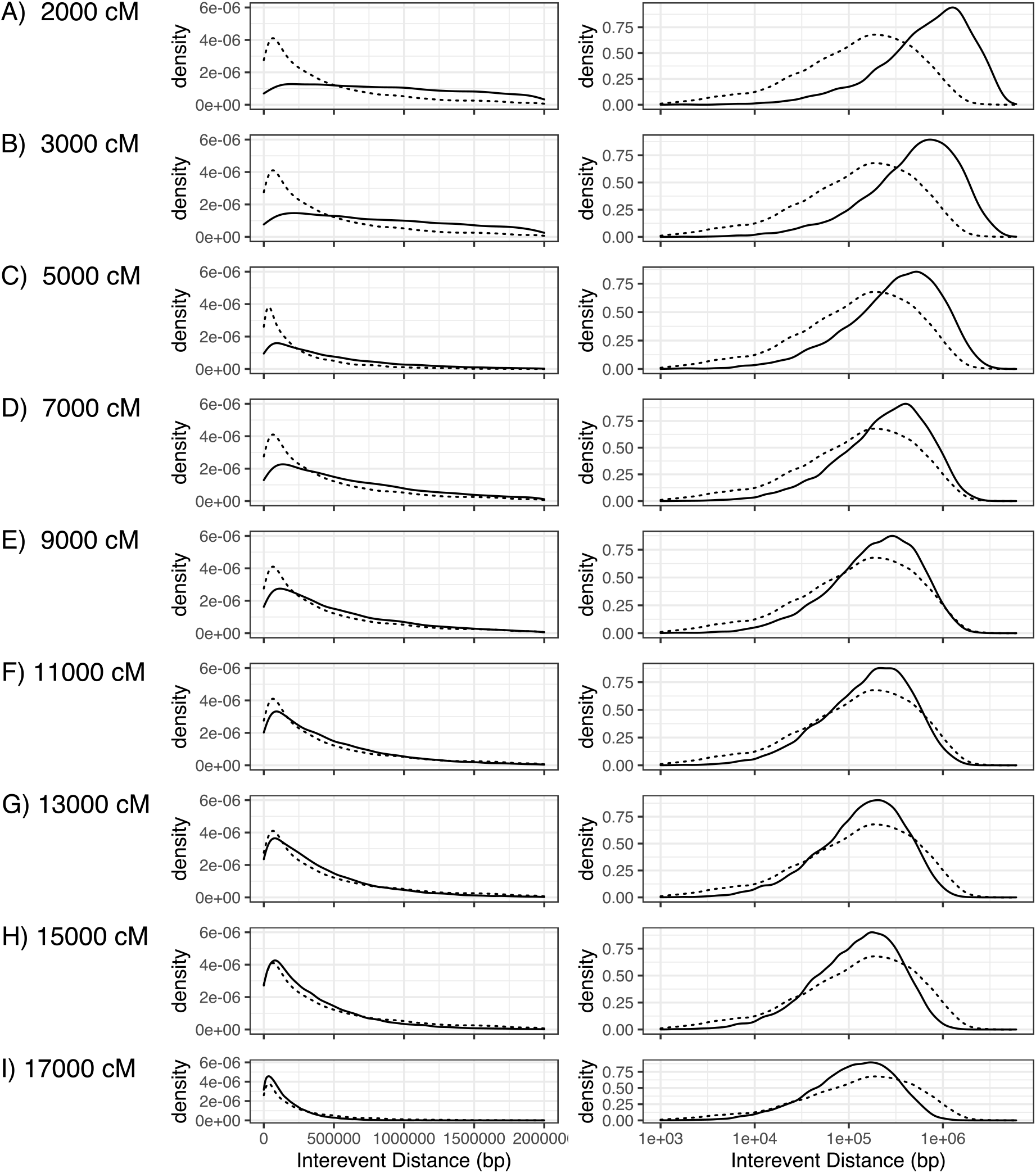
Comparison of cleaned dataset against uniformly distributed simulated maps. **(A-I)** Plots of simulated interevent distances (solid line) versus empirical data (dotted line), right plots show same data as left, but with a log scale on the x-axis.

**Supplemental Fig. 3:**
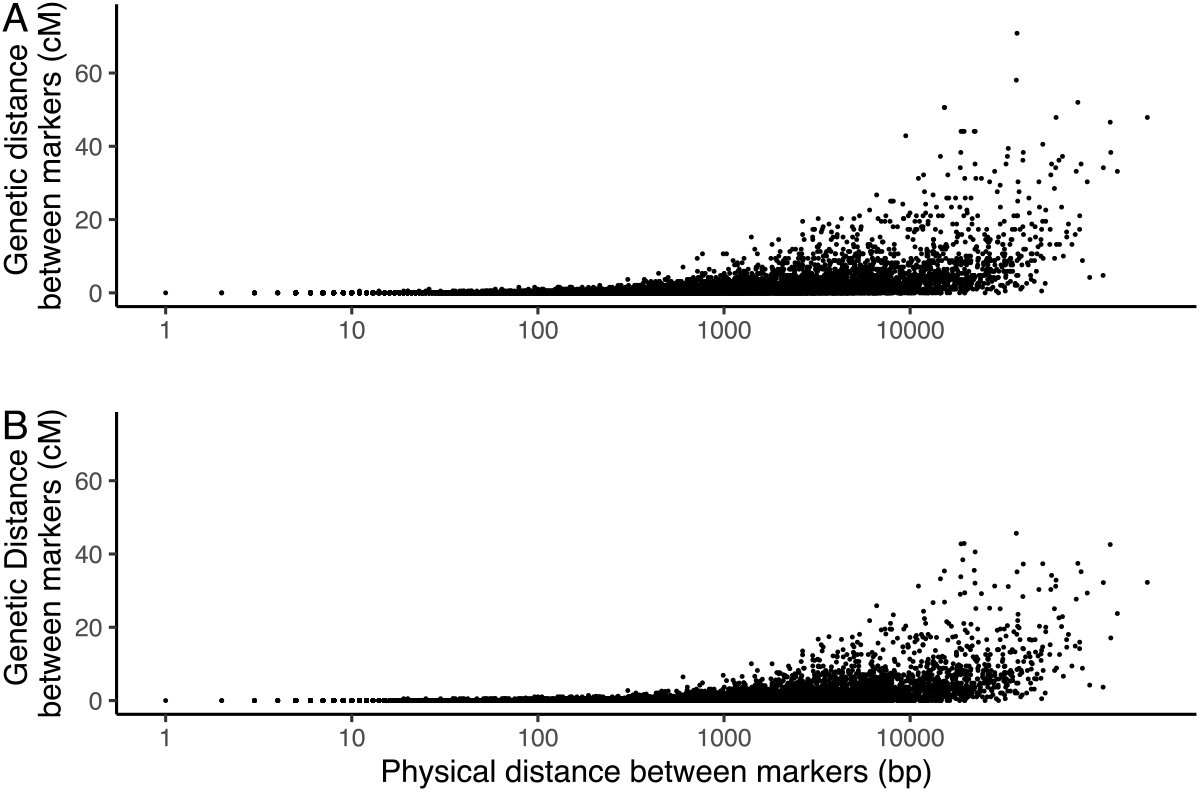
Comparison of physical and genetic distance between markers. **(A)** Comparison of raw dataset, including presumed gene conversions. Each data point indicates the distance between a pair of adjacent markers, with physical distance shown on a log scale. Note that appreciable recombination between markers is only seen above 100 bp. **(B)** Comparison of dataset after removal of gene conversions.

**Supplemental Fig. 4:**
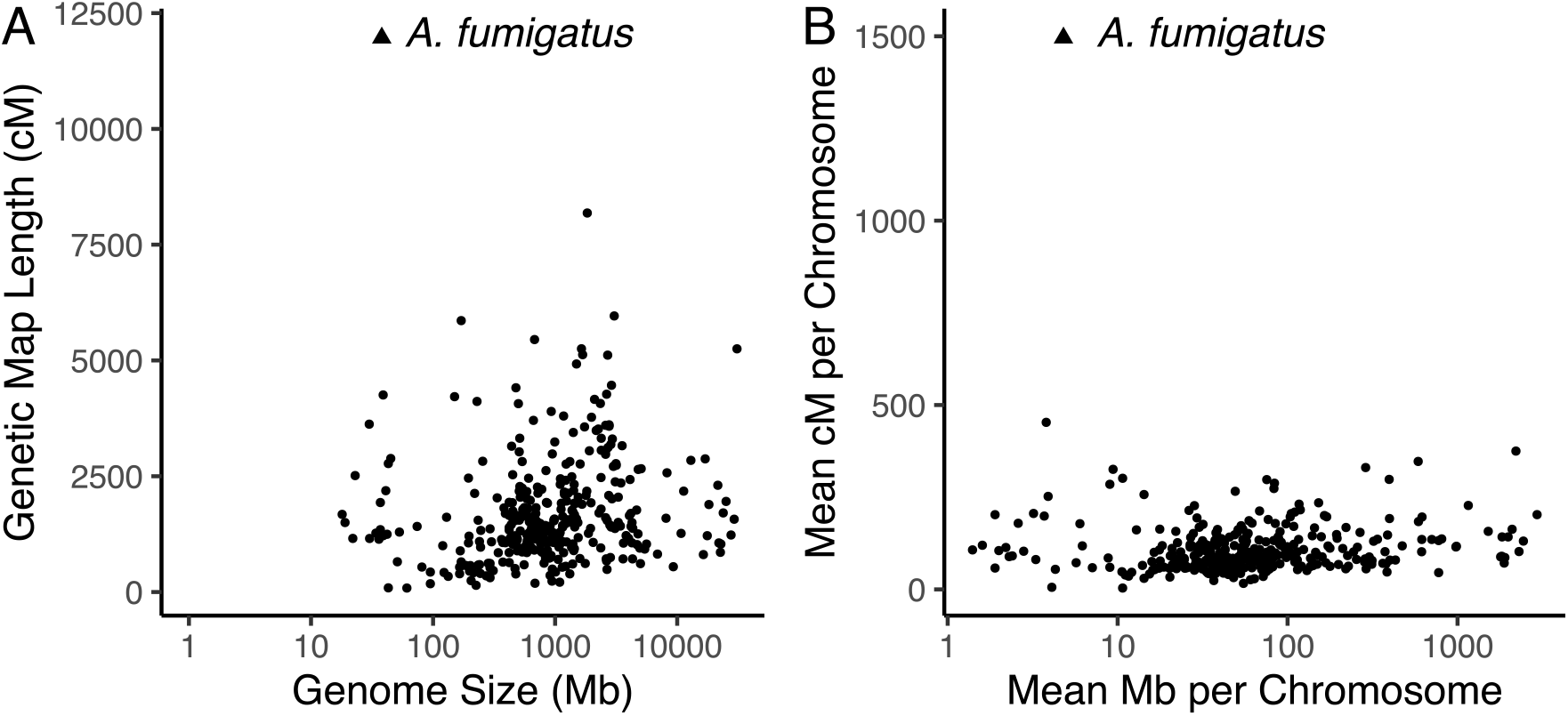
Comparison of genetic map of *A. fumigatus* to Stapley et al. 2017 dataset. **(A)** Small dots indicate values extracted from Stapley et al., 2017, comparing genome size to genetic map length. Larger triangle indicates *A. fumigatus.* **(B)** Similar to A except controlling for chromosome number.

**Supplemental Fig. 5.**
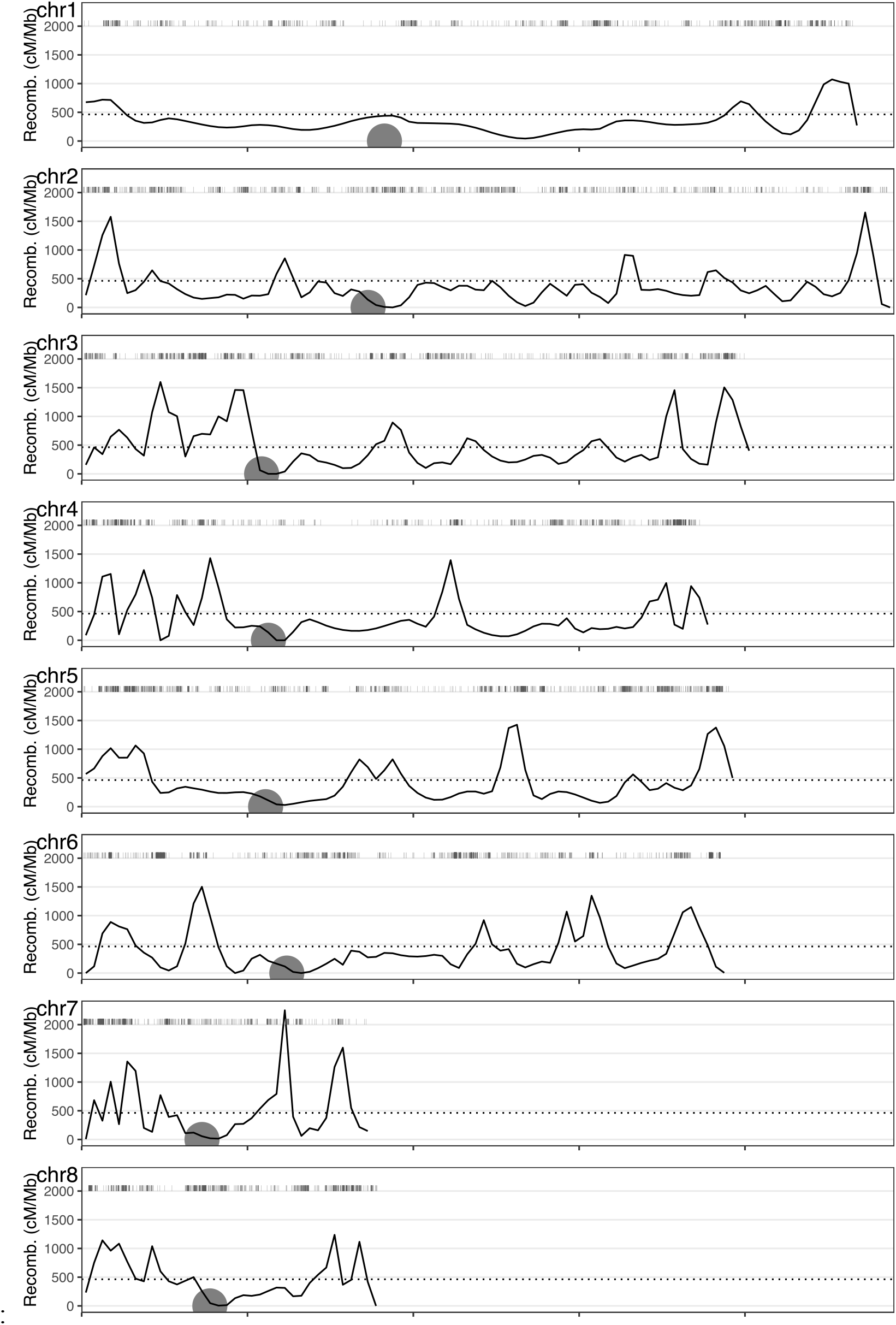
Recombination landscapes of all 8 chromosomes of A. fumigatus. Plots are similar to Figure 2B of main text but shown for all 8 chromosomes. Dotted line indicates the genome-wide average recombination rate, solid line indicates recombination rate across 50kb windows. Small vertical hash lines at top of plots indicate marker positions, and large circles indicate estimated centromere position.

**Supplemental Figure 6:**
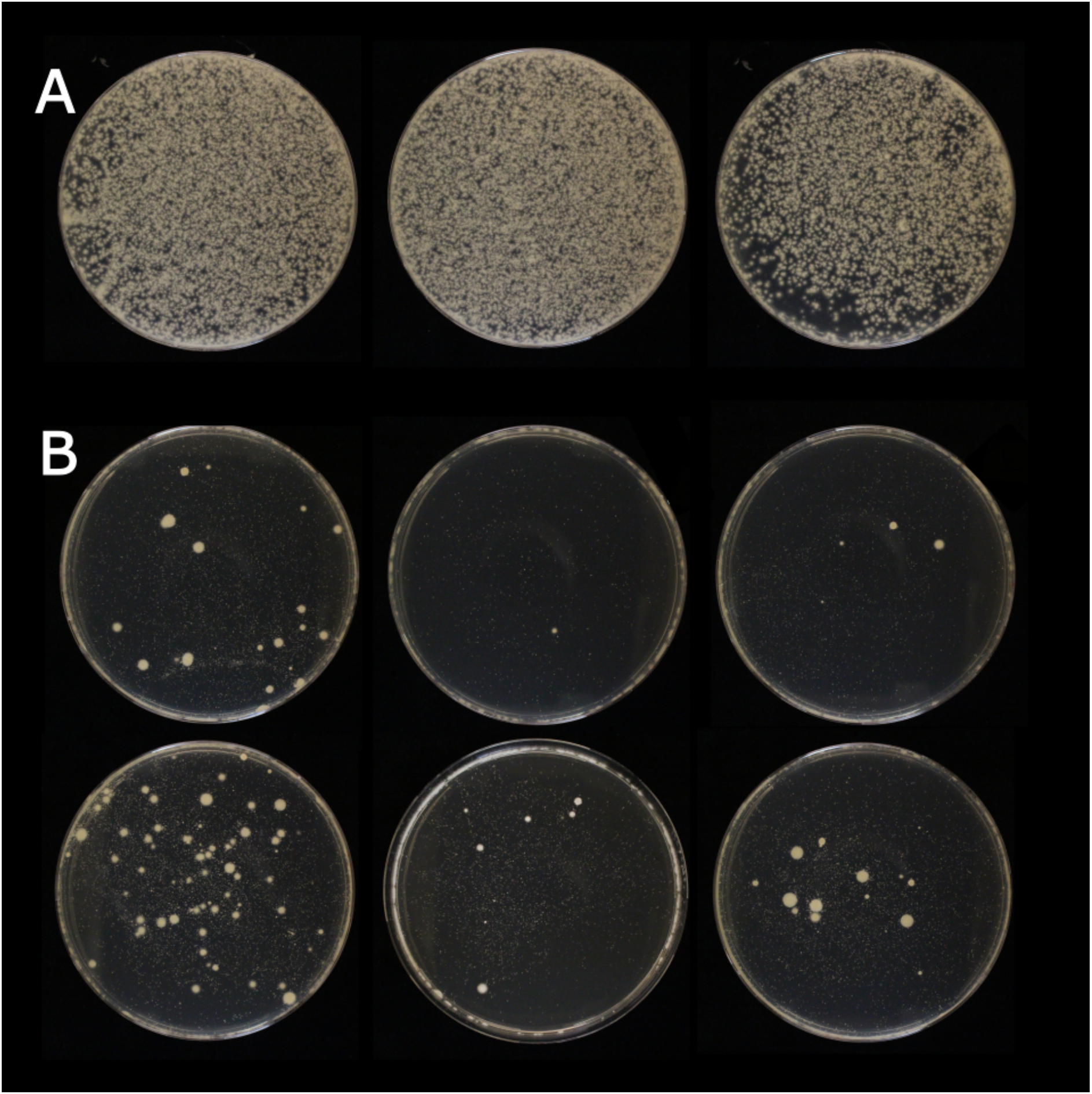
Intragenic *cyp*51A recombination in *A. fumigatus* ascospore progeny. **(A)** Whole contents of single cleistothecia plated on Malt Extract Agar plates. **(B)** Whole contents of single cleistothecia selected for *cyp*51A recombinants by plating instead on Malt Extract Agar + 10μg/mL itraconazole. Bottom right plate is the selection plate used in Fig. 4.

**Supplemental Table 1:**
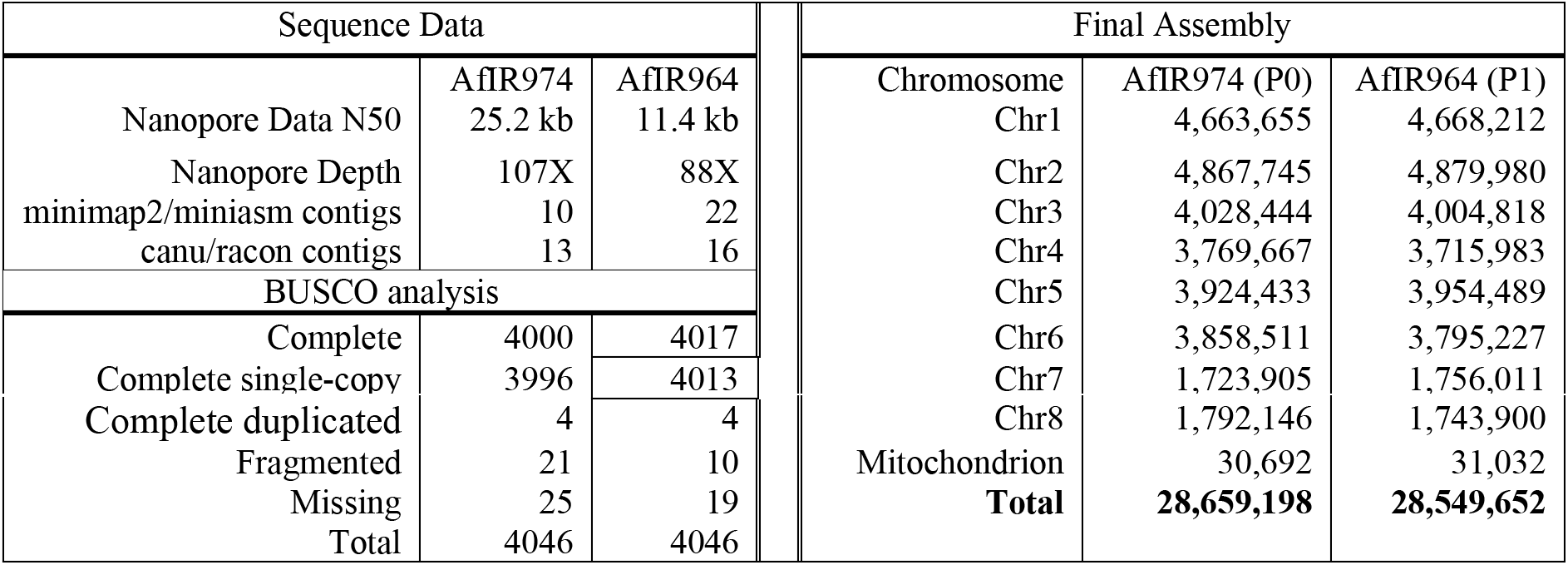
Parental genome assembly statistics.

| Sequence Data |  |  | Final Assembly |  |  |
| --- | --- | --- | --- | --- | --- |
|  | AfIR974 | AfIR964 | Chromosome | AfIR974 (P0) | AfIR964 (P1) |
| Nanopore Data N50 | 25.2 kb | 11.4 kb | Chr1 | 4,663,655 | 4,668,212 |
| Nanopore Depth | 107X | 88X | Chr2 | 4,867,745 | 4,879,980 |
| minimap2/miniasm contigs | 10 | 22 | Chr3 | 4,028,444 | 4,004,818 |
| canu/racon contigs | 13 | 16 | Chr4 | 3,769,667 | 3,715,983 |
| BUSCO analysis |  |  | Chr5 | 3,924,433 | 3,954,489 |
| Complete | 4000 | 4017 | Chr6 | 3,858,511 | 3,795,227 |
| Complete single-copy | 3996 | 4013 | Chr7 | 1,723,905 | 1,756,011 |
| Complete duplicated | 4 | 4 | Chr8 | 1,792,146 | 1,743,900 |
| Fragmented | 21 | 10 | Mitochondrion | 30,692 | 31,032 |
| Missing | 25 | 19 | **Total** | **28,659,198** | **28,549,652** |
| Total | 4046 | 4046 |  |  |  |

**Supplemental Table 2:** Types of variants and their effects.

| Effect of variant | Variant type |  |  |  |
| --- | --- | --- | --- | --- |
|  | MNV | In/del | SNP | sum |
| Non-synonymous | 146 | 67 | 2,212 | 2,425 |
| Intergenic | 829 | 597 | 8,209 | 9,635 |
| Intron | 2 | 20 | 143 | 165 |
| Synonymous | 18 | 0 | 1,870 | 1,888 |
| summed type | 995 | 684 | 12,434 | 14,113 |

**Supplemental Table 3:** Genetic map length statistics. From left to right, chromosome, chromosome length, number of variants detected, recombination events per offspring, raw recombination fraction, rarefied recombination fraction and gene conversion corrected map length is shown.

| Chr | Length (bp) | Number of variants | Mean recombination event / sample | cM recombination fraction | cM rarefied recombination fraction (35 cM) | cM GC corrected |
| --- | --- | --- | --- | --- | --- | --- |
| 1 | 4,663,655 | 1,270 | 16.59 | 1,916 | 1,663 | 1,654 |
| 2 | 4,867,745 | 1,685 | 20.56 | 2,222 | 1,639 | 1,704 |
| 3 | 4,028,444 | 2,061 | 22.51 | 2,509 | 1,952 | 1,887 |
| 4 | 3,769,667 | 2,107 | 16.63 | 1,871 | 1,448 | 1,429 |
| 5 | 3,924,433 | 2,265 | 18.44 | 2,025 | 1,524 | 1,465 |
| 6 | 3,858,511 | 2,015 | 17.64 | 2,058 | 1,685 | 1,615 |
| 7 | 1,723,905 | 1,254 | 9.95 | 1,127 | 929 | 872 |
| 8 | 1,792,146 | 1,456 | 9.87 | 1,115 | 857 | 899 |
| sum | 28,628,506 | 14,113 | 132.18 | 14,848 | 11,698 | 11,525 |

